# Building Foundation Models to Characterize Cellular Interactions via Geometric Self-Supervised Learning on Spatial Genomics

**DOI:** 10.1101/2025.01.25.634867

**Authors:** Yuning You, Zitong Jerry Wang, Kevin Fleisher, Rex Liu, Matt Thomson

## Abstract

Cellular interactions form the fundamental/core circuits that drive development, physiology, and disease within tissues. Advances in spatial genomics (SG) and artificial intelligence (AI) offer unprecedented opportunities to computationally analyze and predict the behavior of cell intricate networks, and to identify interactions that drive disease states. However, challenges arise in both *methodology* and *scalability*: **(i)** how to computationally characterize complicated cellular interactions of multi-scale nature where chemical genes/circuits in individual cells process information and drive interactions among large numbers of diverse cell types, and **(ii)** how to scale up the pipeline to accommodate the increasing volumes of SG data that map transcriptome-scale gene expression and spatial proximity across millions of cells. In this paper, we introduce the **Cellular Interaction Foundation Model** (**CI-FM**), an AI foundation model functioning to analyze and simulate cellular interactions within living tissues. In the CI-FM pipeline, we explicitly capture and embed cellular interactions within microenvironments by leveraging the powerful and scalable geometric graph neural network model, and optimize the characterization of cellular interactions with a novel self-supervised learning objective – we train it to infer gene expressions of cells based upon their interacting microenvironment. As a result, we construct CI-FM with 100 million parameters by consuming SG data of 23 million cells. Our benchmarking experiments show CI-FM effectively infers gene expressions conditional on the microenvironmental contexts: we achieve a high correlation and a low mismatch error (MSE of 1.1% relative to the square median expression), with 79.4% of cells on average being annotated as the similar cell type based on their predicted and actual expressions. We demonstrate the downstream utility of CI-FM by: (i) applying CI-FM to embed tumor samples to capture cellular interactions within tumor microenvironments (ROC-AUC score of 0.76 on classifying sample conditions via linear probing on embeddings), and identifying shared signatures across samples; and (ii) using CI-FM to simulate changes in microenvironmental composition in response to T cell infiltration, which highlights how CI-FM can be leveraged to model cellular responses to tissue perturbations – an essential step toward constructing “AI virtual tissues”. Our model is open source and publicly accessible at https://huggingface.co/ynyou/CIFM.

## Introduction

The cell is the fundamental unit of life, and the cellular communication/interaction establishes the cell-level circuits of living functions – it is indisputably critical for all diseases, from cancer [1] and autoimmune diseases [2] to aging [3] and normal physiology [4]. The scaling of spatial genomics (SG) data [5] and advancements in artificial intelligence (AI) techniques [6] offer unprecedented opportunities to construct computational models to characterize cellular interactions, while current developments do not fully unleash the potential of AI and big SG data: (i) the ineffectiveness of the simplistic model architecture at modeling intricate cellular interaction circuits (e.g. naïve averaging via convolution to represent interactions [7]) which are of multi-scale nature and inherent complexity: in the real-world systems, each cell acts as a node that processes information and changes state, interacting with other nodes through chemical communication, leading to intricate cell interaction networks, where cellular nodes communicate to compute, process information, make decisions, and execute state transitions; and (ii) the challenge in scaling up to accommodate the increasing volumes of SG data that map transcriptome-scale gene expression and spatial proximity across millions of cells, spanning large sections of tissue, organ systems, and disease states [8]. We defer the discussion of more related works to Appdx. A.

In this paper, we aim to build a larger-scale model capable of characterizing cellular interactions to consume massive SG data from varied platforms/sources, in order to function to not only analyze but more importantly, computationally simulate these interactions – an initiative toward constructing “AI virtual tissues”. To this end, we leverage the advanced AI model of geometric graph neural networks (GeoGNNs) [9, 10] – the powerful and scalable model able to explicitly capture and embed interactions within cellular microenvironments (CMEs) via geometric message passing. To optimize the characterization of cell interactions, we develop and implement a novel self-supervised learning pipeline to train GeoGNNs on the vast SG data – around 100 SG samples with 23 million cells and 32 thousand measured genes of four platforms of Visium and Xenium. We refer our model as the cellular interaction foundation model (CI-FM).

Our self-supervised learning pipeline is designed in a masking-reconstruction manner on the SG data of geometric structures. Intuitively, our model is optimized to reconstruct the masked gene expressions of cells based on their microenvironmental contexts. Such a self-supervised learning task is highly intriguing in living systems for the reasons: (i) it requires the model to capture the important cellular interactions within CMEs to deliver the best inference for masked cells, and (ii) the task itself holds significant value in computational simulations of the systems in numerous applications, for instance, it can simulate or “hallucinate” cell state variations in response to tissue-level perturbations in their CMEs.

We benchmark CI-FM and demonstrate that it effectively infers CME context-dependent gene expressions, achieving high correlations and low mismatch errors across varied samples, platforms, and scenarios, including in-sample, cross-sample, and even cross-platform evaluation (e.g., training on Visium and zeroshot evaluation on Xenium). For instance, in the Visium-HD dataset, 79.4% of cells on average are annotated with the top-5 cell typing profile containing shared cell types based on their predicted and actual expressions. Furthermore, we illustrate the downstream utility of CI-FM with two examples. In the first scenario, we use CI-FM embeddings to analyze the CME configurations of different samples. The embeddings well-capture distinguished cellular interactions in healthy and tumor samples, achieving an ROC-AUC score of 0.76 when classifying sample conditions via linear probing. Furthermore, we find that CMEs with highly similar CI-FM embeddings – indicative of shared microenvironmental features across tumor samples – are consistently enriched luminal epithelial cells, suggesting that luminal epithelial cells are core cell types in different tumor types, orchestrating cell-cell communication and maintaining tumor structure. In the second scenario, we apply CI-FM to autoregressively infer CME changes during immune cell infiltration. After the virtual injection of T cells in the breast tumor sample, we observe variations in the populations of different cell types, including epithelial, fibroblast, and endothelial cells.

### CI-FM is trained to reconstruct the gene expressions of masked cells based on their microenvironmental contexts

We hypothesize that the interactions across cells result in the shared information reflected in their gene expressions. Thus, we are intrigued to ask the following question: to what extent can we infer the gene expression of a cell from scratch given the context of its neighbor interacting cells (that constitute CMEs)? To delve into this question, we train a large-scale GeoGNN-based foundation model [9, 10] on massive SG data [5, 11] by optimizing a carefully designed self-supervised learning objective [12, 13]. We refer our model as the cellular interaction foundation model (CI-FM).

The CI-FM workflow consists of three parts: (i) geometric graph featurization of CMEs (Fig. 1B), (ii) masking and encoding/embedding of geometric graphs (Fig. 1C Top), and (iii) padding and decoding/reconstruction of geometric graph features (Fig. 1C Bot). In the foremost step (i), we featurize CMEs in order to feed them into neural networks (Fig. 1B, Appdx. B.1). Considering the *i*th CME (out of *K*, i.e. *i 2* {1, …, *K*}) containing *N*_*i*_ cells with *M*_*i*_ measured genes, the featurized geometric graph is denoted as:

**Fig. 1.**
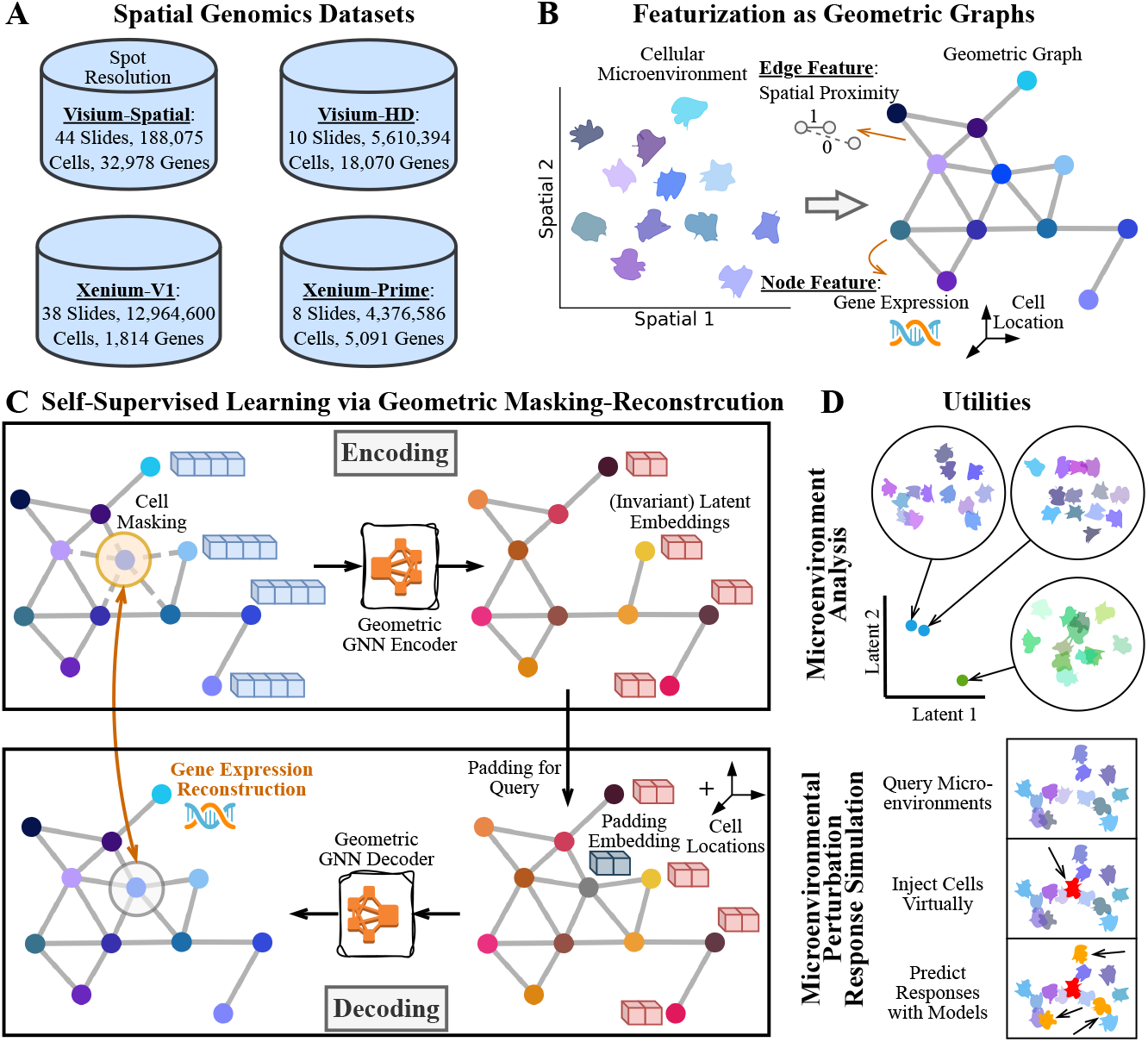
Overview of the CI-FM pipeline. (A) SG data with around 23 million cells of four platforms are curated from the 10x Genomics database. (B) A CME is featurized as a geometric graph with node features of gene expressions and cell locations, and edge features of spatial proximity. (C) CI-FM is trained to reconstruct the gene expressions of masked cells based on their microenvironmental contexts. (D) The utility of CI-FM is demonstrated in the examples of microenvirionment analysis and microenvironmental perturbation perturbation simulation.

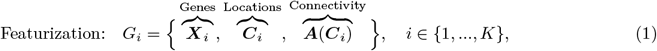

where 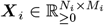 is the node feature matrix of gene expressions, 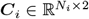 is the coordinate matrix, and 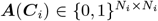 is the adjacency matrix constructed based on the spatial proximity of the coordinates. To collect adequate CMEs for model training, we curate data from the 10x Genomics database [14, 11] (Fig. 1A, Appdx. B.1), a publicly available and extensive resource of spatial genomics data from four different platforms (Visium-Spatial/HD, Xenium-V1/Prime), containing approximately 100 slides of healthy/tumor samples of varied organs, with around 23 million cells and 32 thousand measured genes.

With the featurized geometric graphs, we train CI-FM in a self-supervised manner via masking-reconstruction. In the step (ii), we randomly and uniformly remove nodes within the geometric graph and then use a GeoGNN encoder *f*_enc;*θ*_(·) to embed the remaining structure into the latent space (Fig. 1C Top, Appdx. B.2). In the step (iii), we pad the removed nodes with the learnable padding embeddings ***e***, return them to the geometric graph at their original locations, and then use a GeoGNN decoder *f*_dec;*ϕ*_(·) to reconstruct their masked gene expressions (Fig. 1C Bot, Appdx. B.2). We optimize the GeoGNN encoder/decoder by minimizing the mismatch between the masked and reconstructed gene expressions, formulated as:

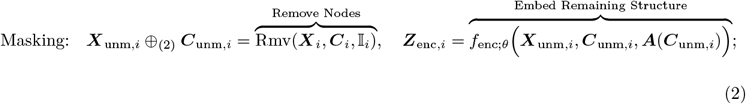

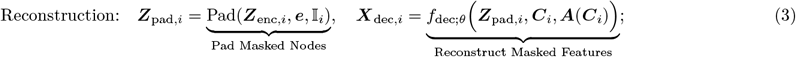

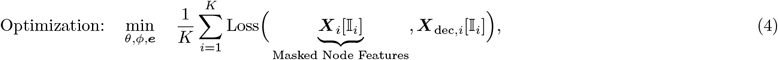

where umn is short for unmasked, ⊕_(2)_ is the concatenation alone the 2nd dimension (feature dimension), Rmv(·), Pad(·) are the removal and padding functions on geometric graphs, respectively, 𝕀_*i*_ is the set of masking indices of the *i*th CME, ***X***[𝕀] denotes matrix indexing, and Loss(·) is the loss function.

We illustrate the utility of CI-FM through two examples, and its capabilities extend beyond then. In the first scenario, we use the embeddings of CI-FM to analyze the CME configurations of tumor samples (Fig. 1D Top, Appdx. B.3). In the second scenario, we apply CI-FM to autoregressively infer changes in the cell states of CMEs after virtually injecting immune cells (Fig. 1D Bot, Appdx. B.3).

### CI-FM effectively infers context-dependent gene expressions across varied spatial genomics samples and platforms

We assess CI-FM by evaluating its precision in inferring the gene expressions of cells, based on the context of their neighboring cells. For benchmarking, we split each slide of SG samples into training, validation, and test regions for model training, hyperparameter tuning, and evaluation, respectively (Fig. 2A). We refer the evaluation with the regional split as in-sample evaluation. We also evaluate using the sample split beyond the regional split, where samples are randomly held out for evaluation (Appdx. C.2), referred as cross-sample evaluation. We compared CI-FM against baselines that include random expressions following parameterized uniform and Bernoulli distributions (with parameters learned from the data), as well as a naive neighborhood average approach that computes the mean expressions of the neighboring cells.

**Fig. 2.**
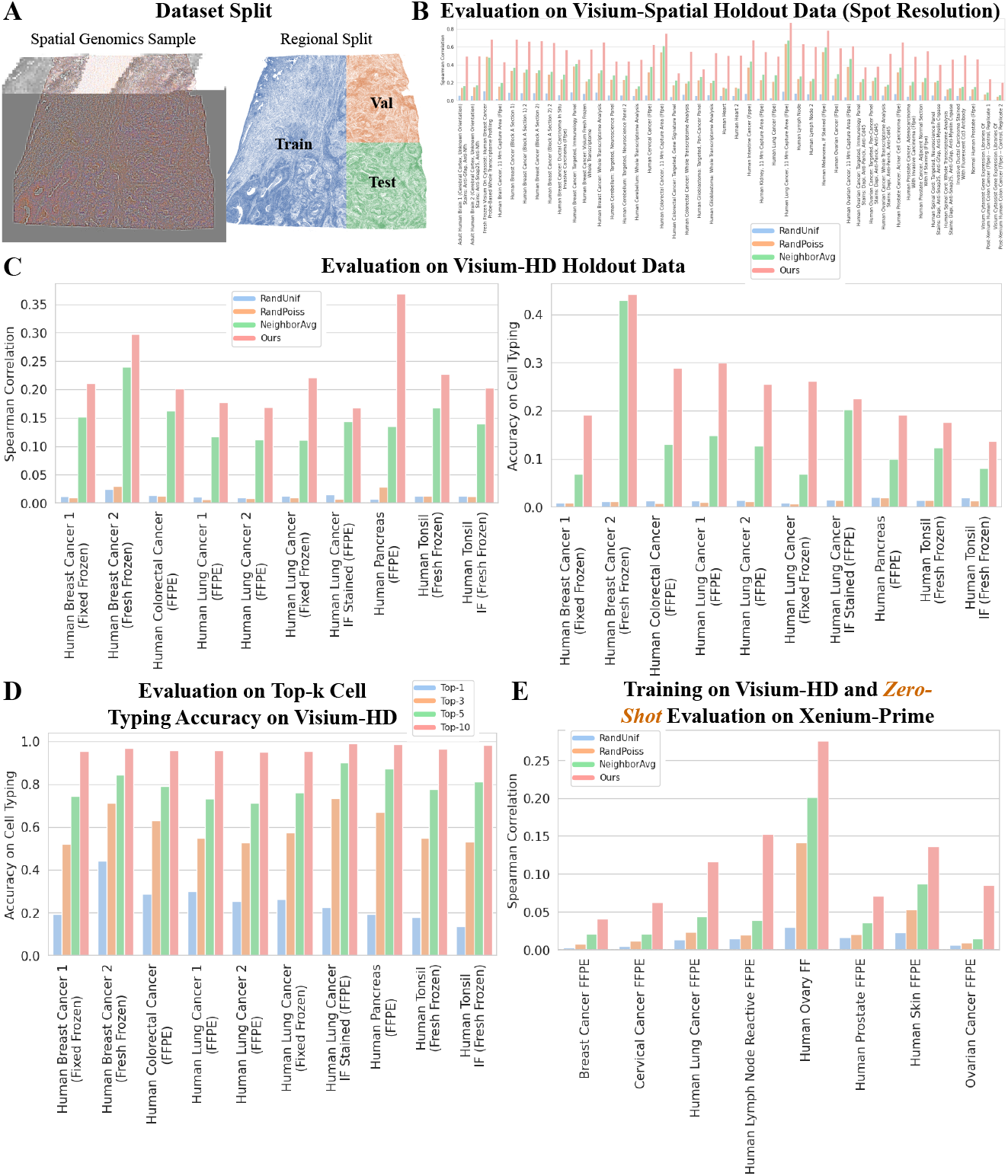
Benchmarking of the CI-FM performance. (A) Dataset split for in-sample evaluation. Cross-sample evaluation and cross-platform evaluation are also conducted. (B) In-sample evaluation on the Visium-Spatial dataset. (C) In-sample evaluation on the Visium-HD dataset. (D) Evaluation on top-k cell typing accuracy of CI-FM on the Visium-HD dataset. (E) Crossplatform evaluation on the Xenium-Prime dataset.

We first evaluate CI-FM on the relatively small-scale Visium-Spatial dataset, which has a lower resolution of spots (Fig. 2B, Appdx. C.1). The evaluation is performed using metrics of correlation and mismatch error: correlation assesses whether the model ranks gene expression correctly, while mismatch error measures how accurately it infers the actual values of gene expression. We observe that CI-FM consistently achieves the best performance across 44 slides in both metrics. We obtain similar observation when evaluating on their 1,000 most differentially expressed genes (Appdx. C.1).

We next evaluate CI-FM on the Visium-HD dataset, which is 50 times larger and offers single-cell resolution (Fig. 2C, Appdx. C.1). In addition to metrics of correlation and mismatch error, we utilize the neural-network based cell typing tool scTab [15] to determine whether reconstructed gene expression is mapped to the same cell type as mapped with the masked gene expression, a measure we term cell typing accuracy. This metric evaluates whether CI-FM infers expressions of the most important genes that is critical in determining cell types. We observe that CI-FM consistently achieves the best performance across 10 slides in all three metrics. We obtain similar observation when evaluating on their 1,000 most differentially expressed genes per sample (Appdx. C.1). Since the cell typing accuracy may be affected by the precision of the cell typing tool itself, we relax the stringency of the cell typing evaluation by considering whether the predicted top-*k* cell types overlap between predictions based on masked and reconstructed gene expressions (Fig. 2D, Appdx. C.1). We observe CI-FM is able to achieve around 80% of accuracy on average when *k* = 5, and nearly 100% accuracy when *k* = 10 (across a total of around 200 cell type labels).

We last assess the transferability of CI-FM by evaluating it on the Xenium-V1 and Xenium-Prime datasets while training on Visium-HD (Fig. 2E), referred as cross-platform evaluation. In spite of the substantial technical differences between Visium (sequencing-based) and Xenium (imaging-based), CI-FM demonstrates effective gene expression inference. In the Xenium-Prime samples, it consistently achieves the highest correlations and lowest mismatch errors, and outperforms in cell typing accuracy for 6 out of 8 slides. However, it underperforms in the Xenium-V1 samples (Appdx. C.3), possibly due to the huge discrepancies in the gene measurement scale: around 17K genes on average in Visium-HD, 300 genes in Xenium-V1, and 5K genes in Xenium-Prime (Fig. 1A). This can be remedied with further finetuning: we further finetune CI-FM on Xenium-V1, which results in the best correlation across all 38 slides (Appdx. C.3).

### CI-FM is able to embed CMEs and characterize microenvironmental patterns across tumor samples

Since the encoding process of CI-FM functions to map the information of CME contexts into latent embeddings, we ask the question that how are the CMEs of different samples distributed within such latent space? We visualize the CME embeddings of Visium and Xenium from CI-FM in a lower-dimensional PCA and UMAP space (Fig. 3A Right). We observe that in the geometries of these two embedding landscape, there existing heterogeneity across tumor samples indicating the sample-specific heterogeneity within CMEs; while there also existing interesting clusters of closely located points indicating shared CMEs across samples.

**Fig. 3.**
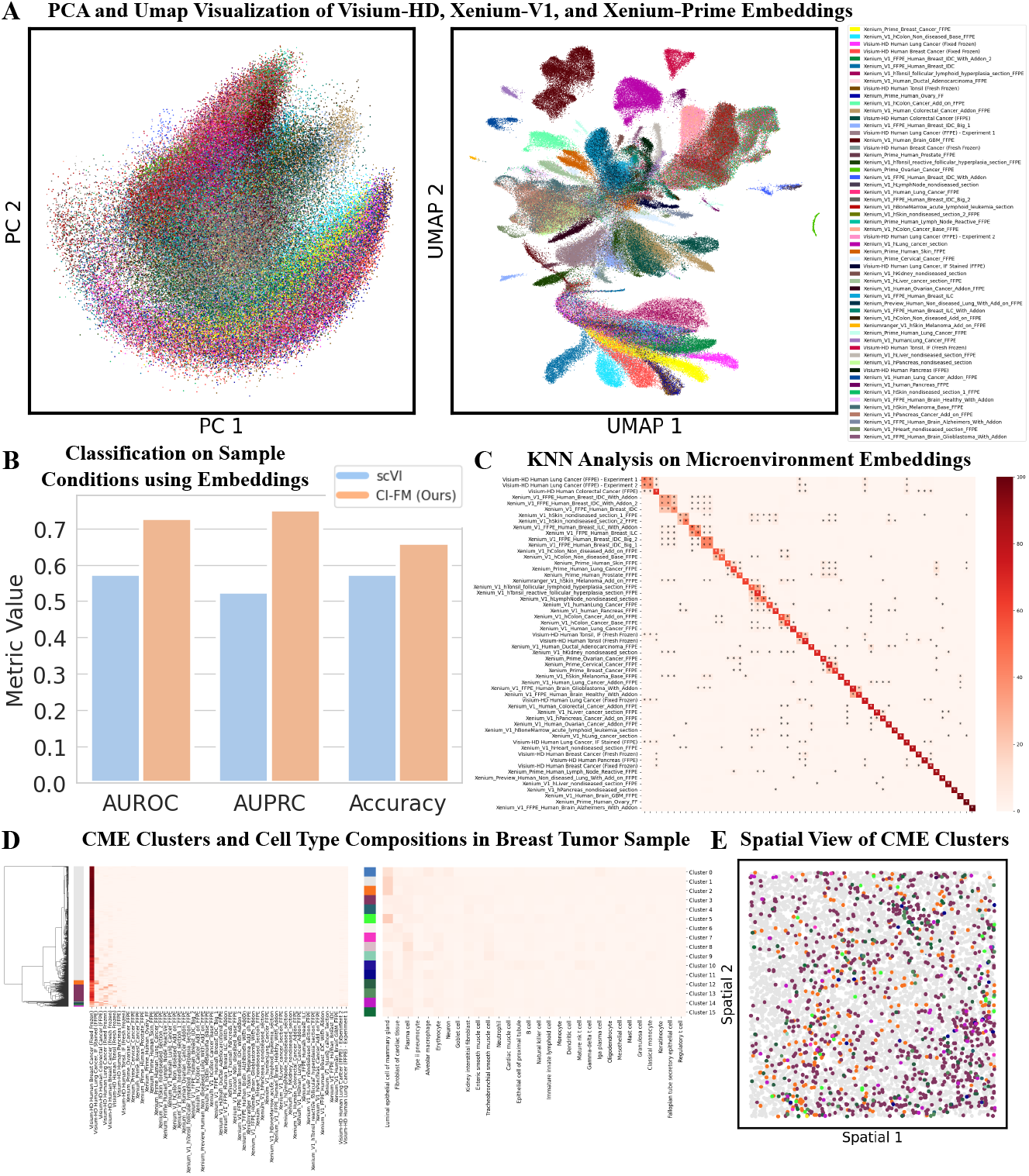
CME embedding with CI-FM. (A) Lower-dimensional visualization of CI-FM embeddings on Visium-HD, Xenium-V1, and Xenium-Prime datasets. (B) Linear probing on CI-FM and scVI embeddings to classify tumor vs non-tumor samples. (C) The fraction (%) of the 100 nearest neighbors queried with CI-FM embeddings across different samples. The value is averaged across all CME embeddings of the entire sample. * denotes a value greater than 1. (D) The KNN profile for each CME in the breast tumor sample without averaging, the hierarchical clustering, and the cell type composition of each cluster. The legend is shared with panel (C). (E) The spatial distribution of each CME cluster.

To quantify how well the microenvironment-level embeddings of CI-FM capture tumor conditions, compared with the cell-level embedding of scVI [16], we perform linear probing to classify tumor and non-tumor samples (Fig. 3B). Specifically, we train a linear classifier on Visium-HD embeddings of tumor and healthy samples and test it on Xenium-Prime sample embeddings. We observe that even this simple linear model achieves decent classification performance in distinguishing tumor from non-tumor samples (ROC-AUC score of 0.76), despite being evaluated on cross-platform data.

To quantify the similarity of CME profiles across samples, we performed a KNN analysis on the CME embeddings, where we ask the question that within the *k* nearest neighbors of each CME embedding, what fraction of these neighbors comes from different samples? As a result, each cell is assigned a KNN fraction vector, where the vector’s length equals the number of samples, its values range from 0 to 100, and its entries sum to 100. We average all the KNN fraction vectors and visualize the sample-level profile (Fig. 3C). Notably, the diagonal blocks capture the shared CMEs within the same tumor types across different samples, and more interestingly the off-diagonal entries capture shared CMEs across different tumor types. We perform the same analysis on the Visium-Spatial dataset and obtain similar observations (Appdx. C.4).

Building upon the above observation, we further investigate a more specific question that what is the cell type composition of these shared CMEs across tumor samples? To answer this question, we retrieve the CME-level KNN profiles (without averaging on samples) and perform clustering (Fig. 3D Left), and then we annotate the cell type composition within each cluster using scTab [15] (Fig. 3D Right). When we cluster the KNN profile of CMEs from a breast tumor sample, we observe that while a large proportion of the CMEs are unique to this particular sample (in gray), there exist a set of CMEs that are appear to be shared across other tumor samples (shown in other colors), including tonsil and colorectal cancer samples. Furthermore, these shared CMEs across tumor samples are enriched with luminal epithelial cells, with their spatial visualization provided (Fig. 3E). We also perform the same analysis and visualization on the other nine samples in the Visium-HD dataset (Appdx. C.4).

### CI-FM is able to simulate CME changes in response to microenvironmental perturbations autoregressively

Since the decoding process of CI-FM maps CME latent embeddings to the potential gene expression profiles associated with CMEs, we ask the question whether we can simulate the evolution of CMEs by iteratively updating their cells? We initialize this process by first virtually injecting cells of interest into CMEs as microenvironmental perturbations, and collect the model outputs that represent the simulated cellular responses to these perturbations. We inject a fixed number of T cells at random locations as perturbations. We visualize the simulation on a breast tumor sample (Fig. 4A). It is important to note that this simulation pipeline is highly general and not limited to specific samples or perturbations.

**Fig. 4.**
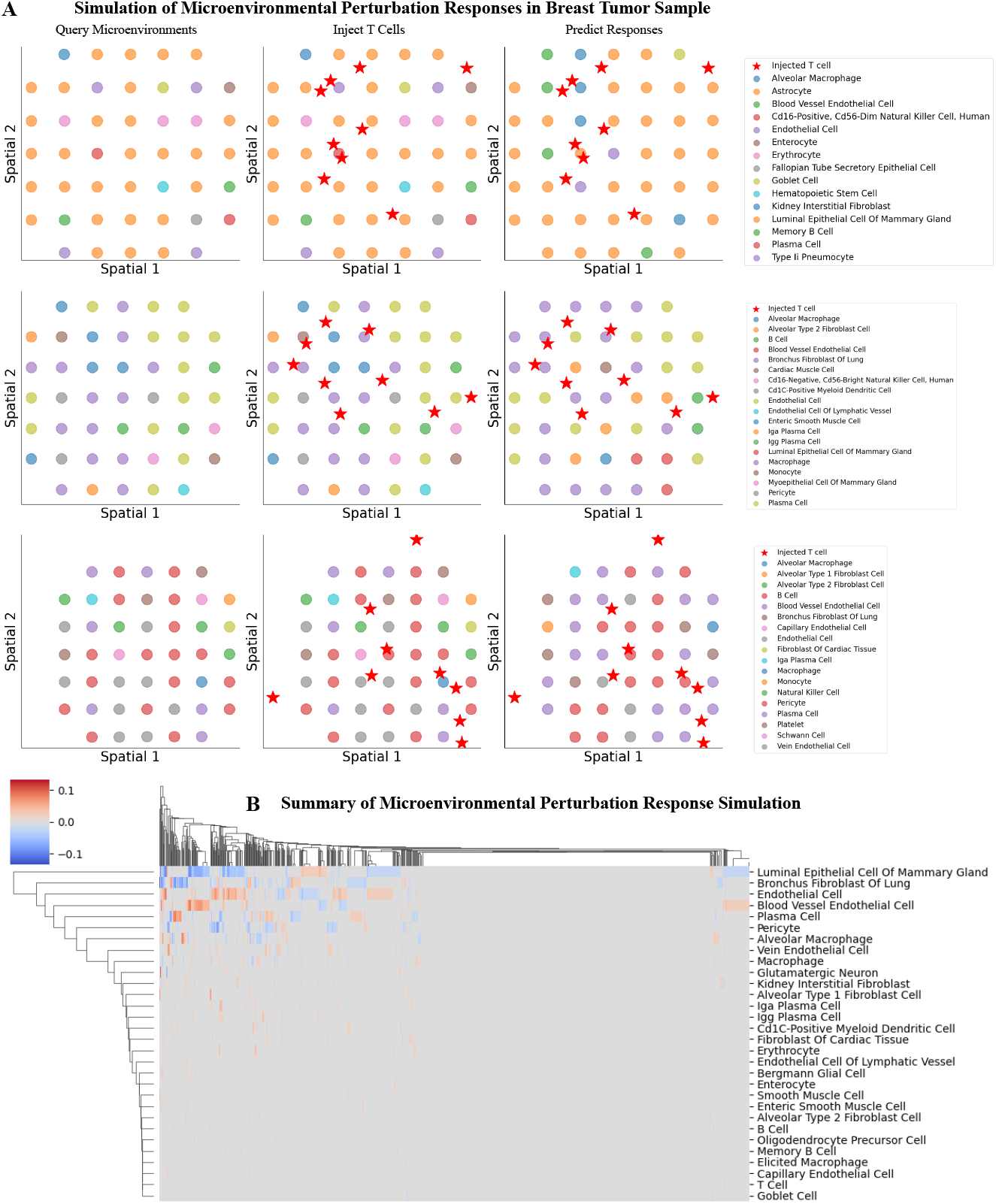
CME response simulation to perturbation with CI-FM. (A) Three examples of the perturbation response simulation in the breast tumor sample. column 1: querying CMEs from samples; column 2: virtually injecting T cells at random locations within CMEs; column 3: masking and predicting gene expressions autoregressively for all cells in the CMEs. (B) Summary of the cell state change in response to the injection of T cells.

We observe a increase in various types of immune cells across the three examples, such as memory B cells. By summarizing changes in cell type composition across a large number CMEs, we observe that CI-FM simulates variations (both increases and decreases) in the populations, in addition to immune cells, of various other cell types including epithelial, fibroblast, and endothelial cells (Fig. 4B). We also simulate and summarize changes in cell type composition across on other nine samples in the Visium-HD dataset (Appdx. C.5).

## Discussion

With the emerging interest in demystifying cellular interactions related to various phenotypes, ranging from aging to cancer, there is a growing demand for computational models designed for this purpose. Advances in artificial intelligence (AI), a powerful data-driven approach, alongside the increasing quality and quantity of single-cell genomic (SG) data, present an exciting opportunity to develop such models. We develop CI-FM here to fully unleash the potential of both modeling and data. Our large-scale model captures interactions across cells and is trained on massive SG datasets. The training process is carefully designed and novel, leveraging cellular interactions to infer missing gene expressions – a feature that enables in silico simulation of living systems. We believe CI-FM represents an important step toward constructing the ultimate biological digital twin, or what could be termed “AI virtual tissue” in the future.

## Appendix A Related Works

### Spatial genomics

Spatial genomics is a rapidly evolving biotechnology that bridges the gap between molecular profiling and spatial context, enabling the study of gene expression within the native tissue architecture. Unlike traditional transcriptomic techniques that analyze dissociated cells and lose spatial information, spatial genomics preserves the spatial relationships between cells, offering unprecedented insights into cellular function, tissue organization, and microenvironmental interactions. Recent technological advances have significantly enhanced our ability to generate multiplexed tissue imaging data that captures cellular neighborhoods within intact tissues. A wide range of platforms, including sequencing-based methods (e.g., Visium [14], Stereo-seq [17]) and imaging-based approaches (e.g., Xenium, seqFISH [5], MERFISH [18]), now allow for the simultaneous measurement of thousands of genes with high spatial precision. Moreover, the rapid development of new platforms and methods continues to increase the diversity and complexity of spatial genomics data, enabling deeper insights into different biological conditions.

#### AI virtual tissue

AI-powered virtual tissue models, a brand-new concept, represent an exciting intersection of artificial intelligence and biological research, enabling the analysis of tissue structure and function with unprecedented precision [6]. These models leverage advanced machine learning algorithms to analyze tissue-level data such as spatial genomics, transcriptomics, proteomics, and imaging. Compared to recent concurrent works with a similar perspective [19, 20] that are based on transformer architectures, our CI-FM demonstrates superior scalability due to the unique advantages of geometric graph neural networks.

#### Geometric graph neural networks

Geometric graph neural networks (GeoGNNs) are an extension of traditional graph neural networks (GNNs) [21, 22], designed to process data that resides on geometric structures like manifolds, point clouds, or other spatially embedded graphs, enabling superior performance in various domains such as computational biology, physics simulations, robotics, and computer vision [23, 9]. Compared to GNNs, GeoGNNs have the advantage of better expressiveness. They leverage the geometric and topological properties of data [10]. Compared to transformers [24, 25], GeoGNNs are more scalable, as they employ local message passing rather than global (fully connected) mechanisms.

#### (Geometric) graph self-supervised learning

Self-supervised learning on graphs is shown to learn more generalizable, transferable and robust graph representations, through exploiting vast unlabelled data [26, 27]. The main advantage of self-supervised learning is that it does not require any human annotation which is usually resource-intensive in biology, enabling the utilization of rich data resources to their fullest potential [13, 12]. Recent efforts have also extended this technique to geometric graphs, demonstrating its effectiveness in various applications, such as molecular representation learning [28, 29].

## Appendix B Extended Methodological Details

### B.1 Spatial Genomics Data Preprocessing and Geometric Graph Featurization

We download the raw count SG data of Visium and Xenium from 10x Genomics (https://www.10xgenomics.com/datasets), using filters “Visium Spatial” and “Xenium In Situ” under Platform, and “Human” under Species. We read the raw counts with coordinates measured in micrometers, and their metadata into the AnnData format [30]. We remove mitochondrial genes, void cells (those without detected gene expression), and retain only cells annotated as “in tissue” based on their histological images, and then normalize gene counts and conduct log1p-transformation. For Visium-HD data, we select a bin size of 8 × 8*μ*m as recommended. We eventually obtain 100 slides, with 23,139,655 cells and 32,986 measured genes (Fig. B1A).

For benchmarking our approach, we split each slide into training, validation, and test regions by segmenting it. We define the segmentation rule as follows: using a slide-specific threshold along the *x*- and *y*-axes as *x*_thres_, *y*_thres_, we allocate a cell with coordinates (*x, y*) to the training data if *x* ≤ *x*_thres_; to the validation data if *x* > *x*_thres_ and *y* ≥ *y*_thres_; and to the test data if *x* > *x*_thres_ and *y* < *y*_thres_. This unbiased split allows us to preserve the CMEs of each cell as much as possible. We compute the thresholds as follows: given the minimum and maximum coordinates of each slide along the *x*- and *y*-axes as *x*_min_, *x*_max_, *y*_min_, *y*_max_, we calculate *x*_thres_ = *x*_min_ + (*x*_max_ − *x*_min_) × 0.6, and *y*_thres_ = *y*_min_ + (*y*_max_ − *y*_min_) × 0.5. The number of cells in training, validation, and testing regions follows an approximate 3:1:1 ratio in each slide (Fig. B1B).

With the above preprocessing, we obtain multiple slides of SG data containing information about the expressions and locations of cells. For the *i*th slide containing *N*_*i*_ cells and *M*_*i*_ measured genes, we featurize it into a geometric graph 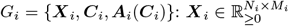, the node feature matrix, is derived from gene expressions; 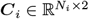 the geometric feature matrix, is derived from spatial locations; and 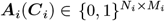, the adjacency matrix, is constructed from ***C***_*i*_ to capture spatial proximity information. We build a radius graph for the adjacency matrix that given the radius threshold *r*_thres_, the adjacency matrix is computed as: 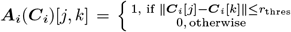, with the radius threshold *r*_thres_ set to 20*μ*m for Visium-HD, Xenium-V1 and Xenium-Prime and 150*μ*m for Visium-Spatial of spot resolution.

**Fig. B1].**
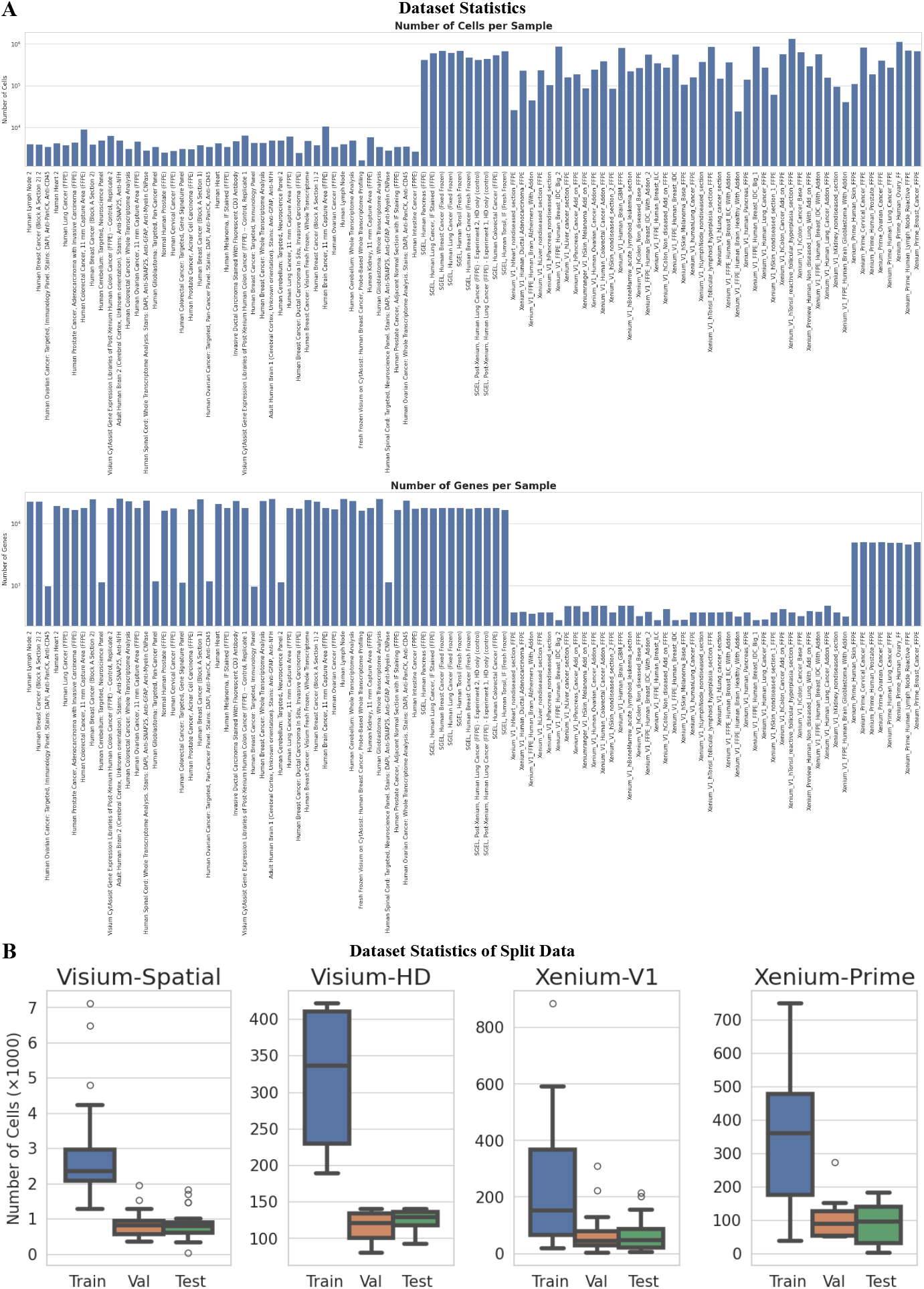
Dataset statistics. (A) Number of cells and measured genes per sample in the Visium and Xenium datasets. (B) Number of cells in the training, validation and test in the regional split.

### B.2 Geometric Graph Neural Network Encoder and Decoder JHUH

GeoGNNs are capable of processing geometric data while respecting their inherent symmetry of permutation and spatial transformations [9, 10], which is important for cellular systems as biological functions remain unaffected by permutations in cell features or by global transformations (of rotation and translation) in location. Specifically, a GeoGNN *f* (·) takes a geometric graph as input and outputs a (*D* + 2)-dimensional invariant and equivariant embedding for each node (cell) as ***Z***_inv_ ⊕_(2)_ ***Z***_eqv_ = *f* (***X, C, A***(***C***)), where ⊕_(2)_ is the concatenation alone the 2nd dimension (feature dimension), ***Z***_inv_ ∈ ℝ^*N*×*D*^ is the invariant embedding matrix, and ***Z***_eqv_ ∈ ℝ^*N*×2^ the equivariant embedding matrix. It respects the symmetry via strictly satisfying:

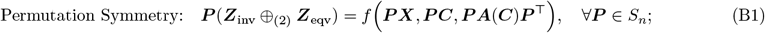

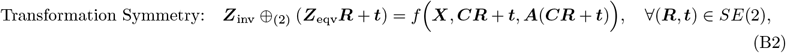

where *S*_*n*_ is the group of permutation of *n* elements, and *SE*(2) is the Special Euclidean group in 2D.

We leverage the E(n) equivariant graph neural network (EGNN) [9] as the base GeoGNN encoder and decoder architecture for its effectiveness and efficiency. Considering the input and output of the *l*th EGNN layer (out of *L, l* ∈{1, …, *L*}) is denoted as 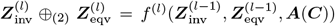 where 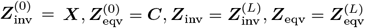, the layer-wise message passing of the *i*th node is formulated as:

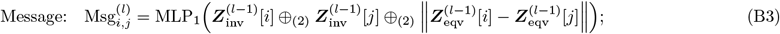

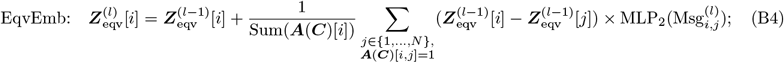

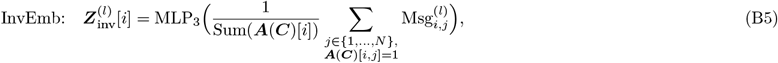

where MLP(·) denotes the multilayer perceptron, Sum(·) represents the summation function, and the computation can be parallelized across all nodes simultaneously. We rely on the final invariant representation ***Z***_inv_ for encoder embeddings and decoder reconstructions, so the final layer I/O is formulated as 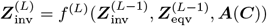.

One issue with the current GeoGNN architectures is the presence of void node biases, where the addition of dummy void nodes unnecessarily affects the representations of real nodes. In cellular systems, the addition of non-realistic void cells (e.g. without any gene being expressed) introduces bias into GeoGNN models, affecting cell representation in arbitrary directions. Specifically, we aim to further enforce the GeoGNN model to respect the void symmetry by strictly satisfying:

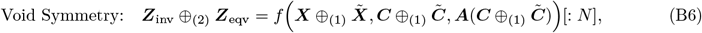

where 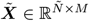 represents the problem-specific void node features, 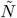 is the number of void nodes, and 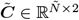 denotes the arbitrary locations of the void nodes. We focus on eliminating the bias for void cells, specifically when 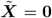. We achieve this by proposing a modified void-invariant architecture based upon EGNNs (Eqs. (B3) - (B5)), with message-passing formulated as:

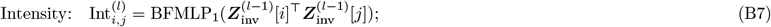

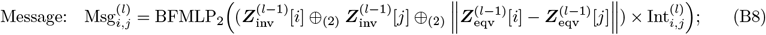

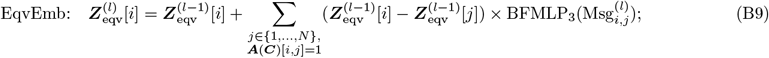

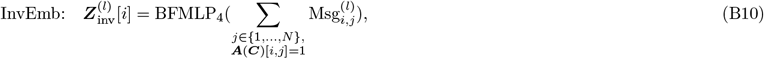

where BFMLP(·) denotes the bias-free multilayer perceptron containing no bias weights.

The key components introduced to guarantee void symmetry (Eq. (B6)) are as follows: 1) the introduction of the intensity term (Eq. (B7)) ensuring that the zero intensity if either the *i*th or *j*th node is void, thereby nullifying the void message; 2) the employment of bias-free multilayer perceptron to preserve the void message; 3) the utilization of sum pooling (rather than mean pooling) to eliminate the population bias in representation resulting from void nodes. More importantly, the void-invariant architecture allows the computation of derivatives at void features, i.e. 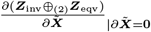 exists, which is highly valuable for numerous downstream applications requiring gradient information (e.g. counterfactual search [8]).

### B.3 Geometric Self-Supervised Learning

We train the GeoGNN in a self-supervised manner through a masking-reconstruction approach. In the encoding stage, we randomly remove 5% of the nodes for masking. Specifically, given an input geometric graph *G* = {***X, C, A***(***C***)}, and denoting the masked and unmasked indices as 𝕀 and 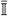, respectively, the masking process is performed by removing the nodes as ***X***_unm_ ⊕_(2)_ ***C***_unm_ = Rmv(***X, C***, 𝕀) where 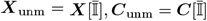. A GeoGNN encoder *f*_enc;*θ*_ (·) parametrized with *θ* is then applied to embed the geometric graph as ***Z***_enc_ = *f*_enc;*θ*_ (***X***_unm_, ***C***_unm_, ***A***(***C***_unm_)). In the decoding stage, we pad the masked node in the latent space with a learnable padding embedding ***e*** ∈ ℝ^*D*^. Specifically, the padding process is performed as ***Z***_pad_ = Pad(***Z***_enc_, ***e***, 𝕀) where ***Z***_pad_[*i*]= ***e***, ∀*i* ∈ 𝕀. A GeoGNN decoder *f*_dec;*ϕ*_(·) parametrized with is then applied to reconstruct the masked node features as ***X***_dec_ = *f*_dec;*ϕ*_(***Z***_pad_, ***C, A***(***C***)).

We optimize the model by minimizing the discrepancy between the masked and reconstructed gene expressions. We optimize on a balanced MSE loss on all the *K* samples with the optimization formulated as:

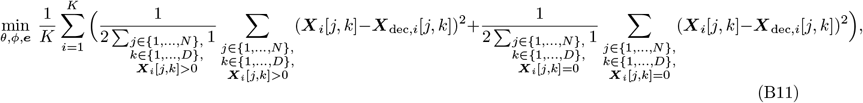

where the subscript represents the sample index.

## Appendix C Extended Results

### C.1 Evaluation on Alternative Metrics

The metrics used for correlation are Spearman correlation and for mismatch error the mean square error (MSE). We further assess CI-FM on the balanced MSE metric expressed as 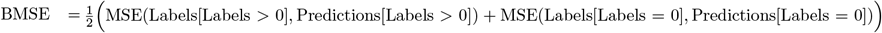 to mitigate the dominance of non-expressed genes, and on Spearman correlation for non-zero-expressing genes to evaluate the model’s ranking accuracy for expressed genes. Moreover, we perform the same evaluation process on the 1,000 most differentially expressed genes per sample computed with Scanpy [30]. We evaluate these metrics on the Visium-Spatial dataset (Fig. C2) and the Visium-HD dataset (Fig. C3).

The cell typing accuracy, computed by mapping reconstructed gene expression to the same cell type as mapped using masked gene expression, may be affected by the precision of the cell typing tool itself. To address this, we relax the stringency of the cell typing evaluation by considering whether the predicted top-*k* cell types overlap between predictions based on masked and reconstructed gene expressions. We evaluate this with *k* set to 1 (original evaluation), 3, 5, and 10. The top-*k* cell typing accuracy is computed on the Xenium-V1 and Xenium-Prime datasets (Fig. C4).

### C.2 Evaluation with Cross-Sample Split

We perform a cross-sample split by randomly holding out samples for testing and using a regional split of the remaining data for training and validation. We evaluate CI-FM on the Visium-Spatial dataset, by holding out 9 of the 44 samples for test and using the same metrics of correlation and mismatch error (Fig. C5).

### C.3 Evaluation on Cross-Platform Spatial Genomics Data

We perform a cross-platform evaluation by training CI-FM on the Visium-HD dataset and evaluate on the Xenium dataset. To ensure that the input gene expression features maintain the same semantics when fed into the model, we perform channel matching before evaluation: we match gene channels across platforms if they are measured on both, and zero-initialize them otherwise. In our datasets, nearly all gene channels are matched between Visium-HD and Xenium, with only a few exceptions. We perform zero-shot evaluation and finetuning evaluation: in zero-shot evaluation, we directly assess the model trained on Visium-HD after channel matching, and in finetuning evaluation, we perform one epoch of finetuning before the evaluation. We evaluate on both the Xenium-V1 and Xenium-Prime datasets (Fig. C6).

**Fig. C2].**
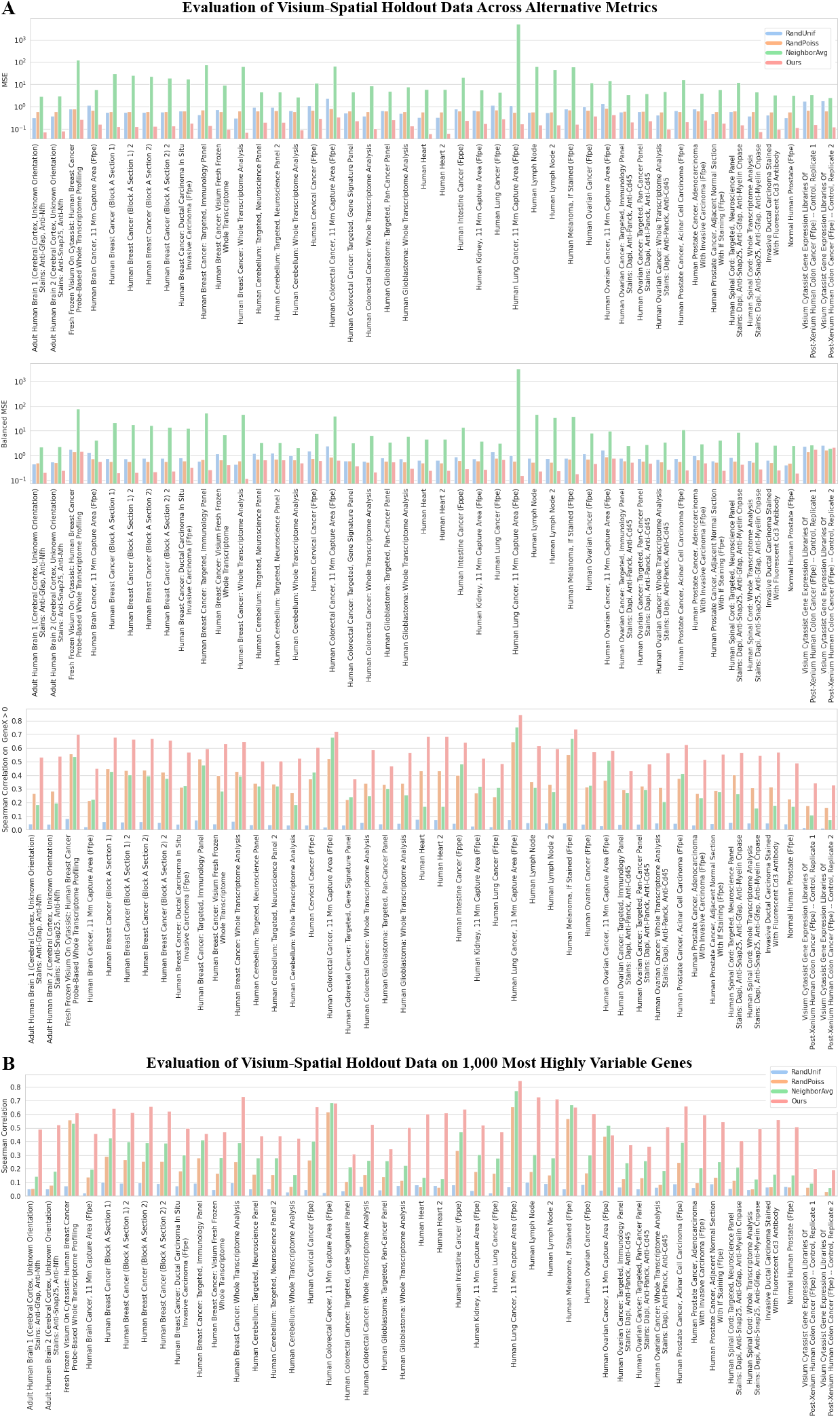
Extended benchmarking results on the Visium-Spatial dataset. (A) Evaluation results on MSE, balanced MSE and Spearman correlation on non-zero-expressing genes. (B) Evaluation results on the 1,000 most differentially expressed genes.

### C.4 Extended Embedding Analysis Results

We further investigate the question that what is the cell type composition of the CMEs across different samples. We pursue this in a manner similar to that described in the main text, applied to all other nine samples in the Visium-HD dataset: CME clustering, cell type annotation, and spatial distribution visualization (Fig. C8, Fig. C9). Additionally, we perform similar lower-dimensional embedding visualizations and KNN analyses for the samples in the Visium-Spatial dataset (Fig. C10).

**Fig. C3].**
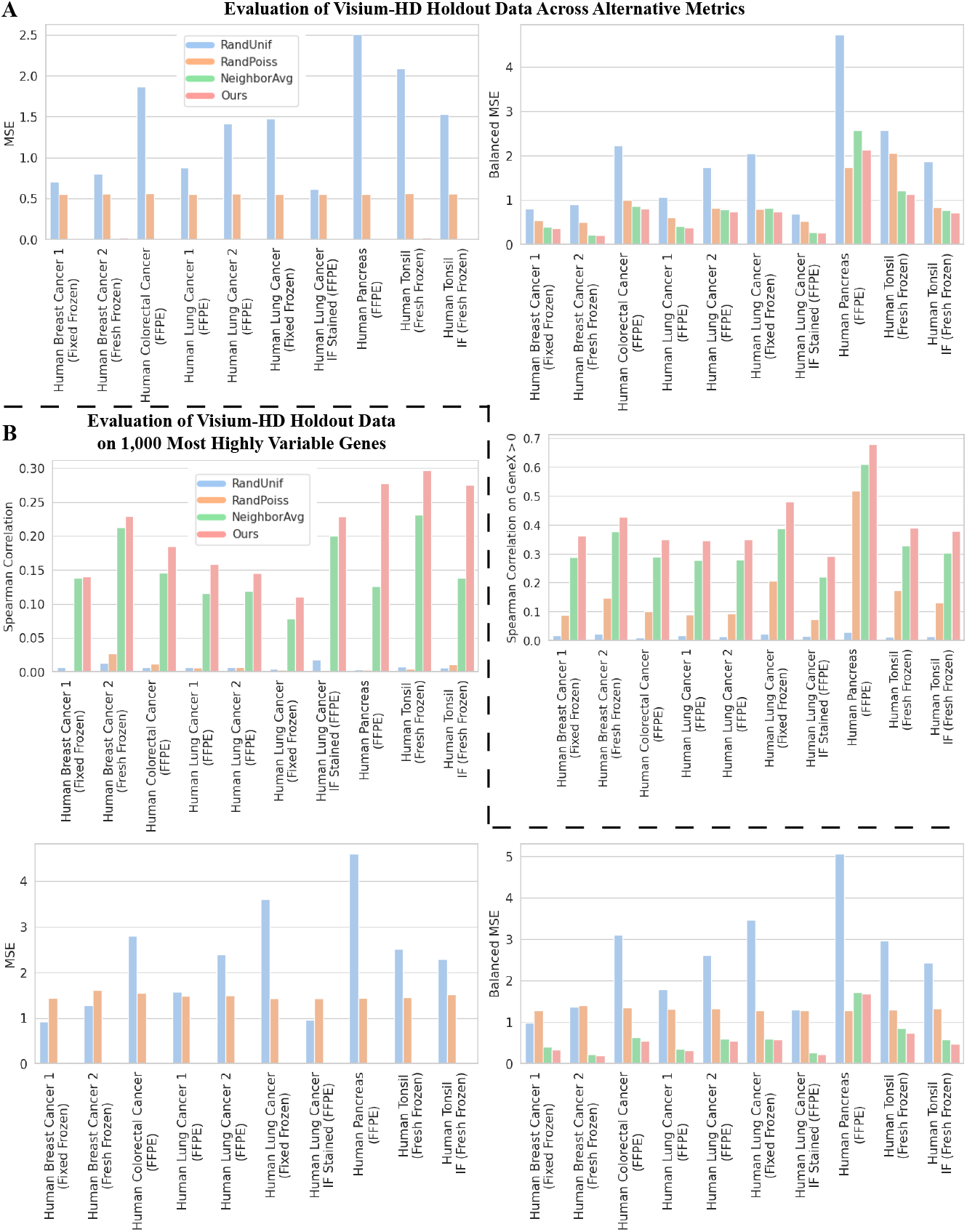
Extended benchmarking results on the Visium-HD dataset. (A) Evaluation results on MSE, balanced MSE and Spearman correlation on non-zero-expressing genes. (B) Evaluation results on the 1,000 most differentially expressed genes.

### C.5 Extended Perturbation Response Simulation Results

We further simulate and summarize changes in cell type composition across a large number CMEs on other nine samples in the Visium-HD dataset (Fig. C11).

**Fig. C4].**
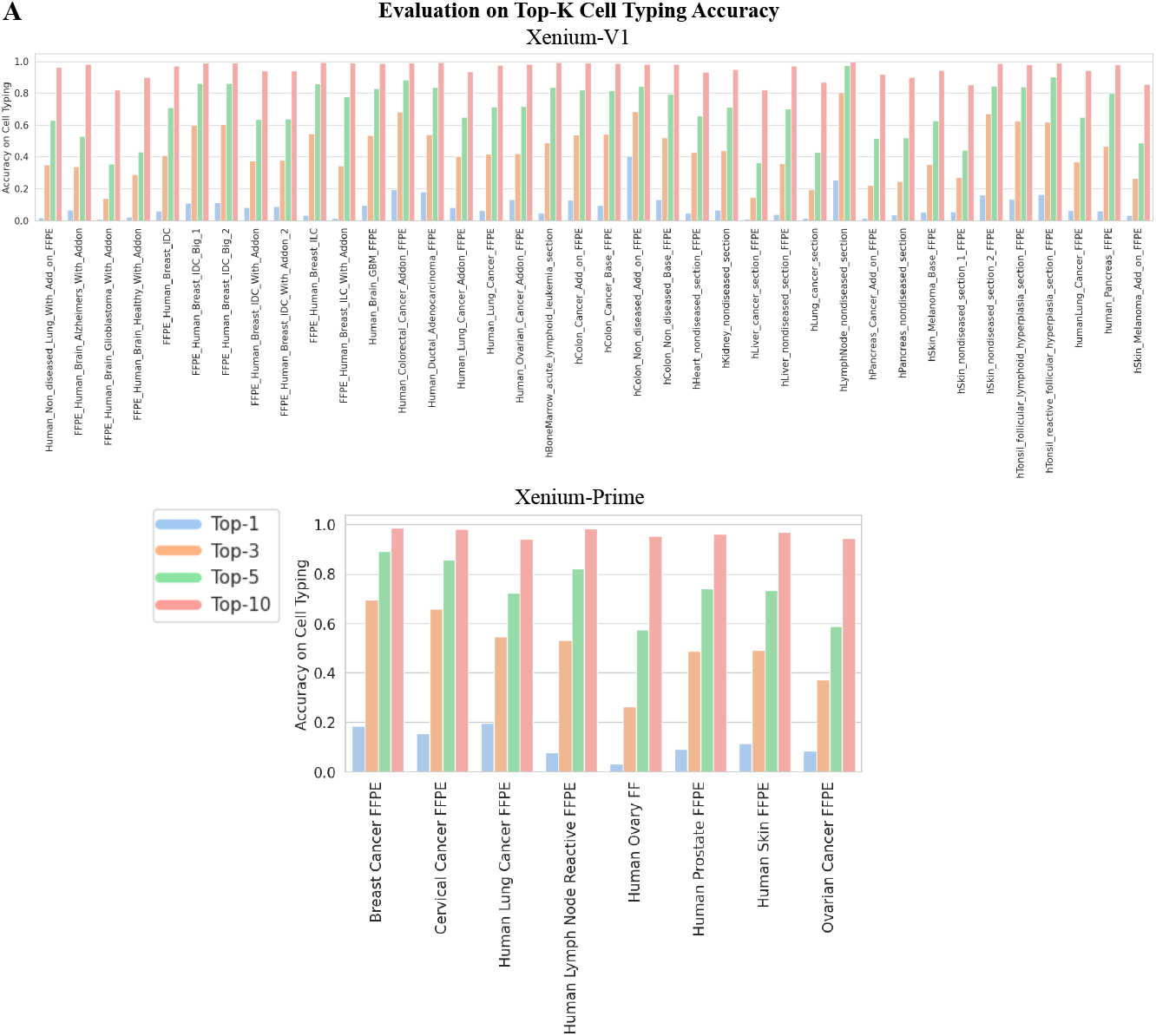
Evaluation on top-*k* cell typing accuracy. (A) Evaluation results on the Visium-HD, Xenium-V1 and Xenium-Prime datasets.

**Fig. C5].**
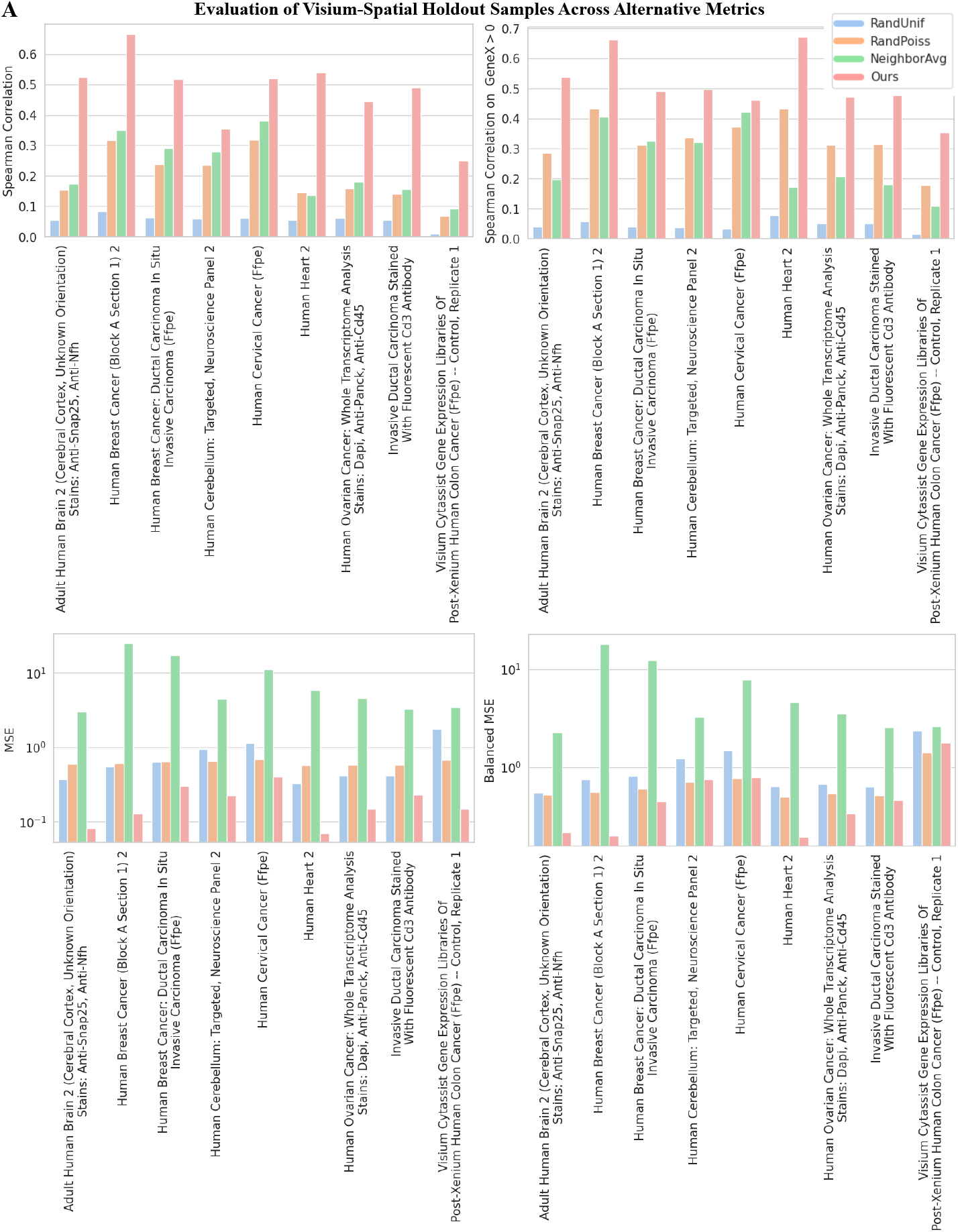
Cross-sample evaluation on the Visium-Spatial dataset. (A) Evaluation results on Spearman correlation, Spearman correlation on non-zero-expressing genes MSE and balanced MSE.

**Fig. C6].**
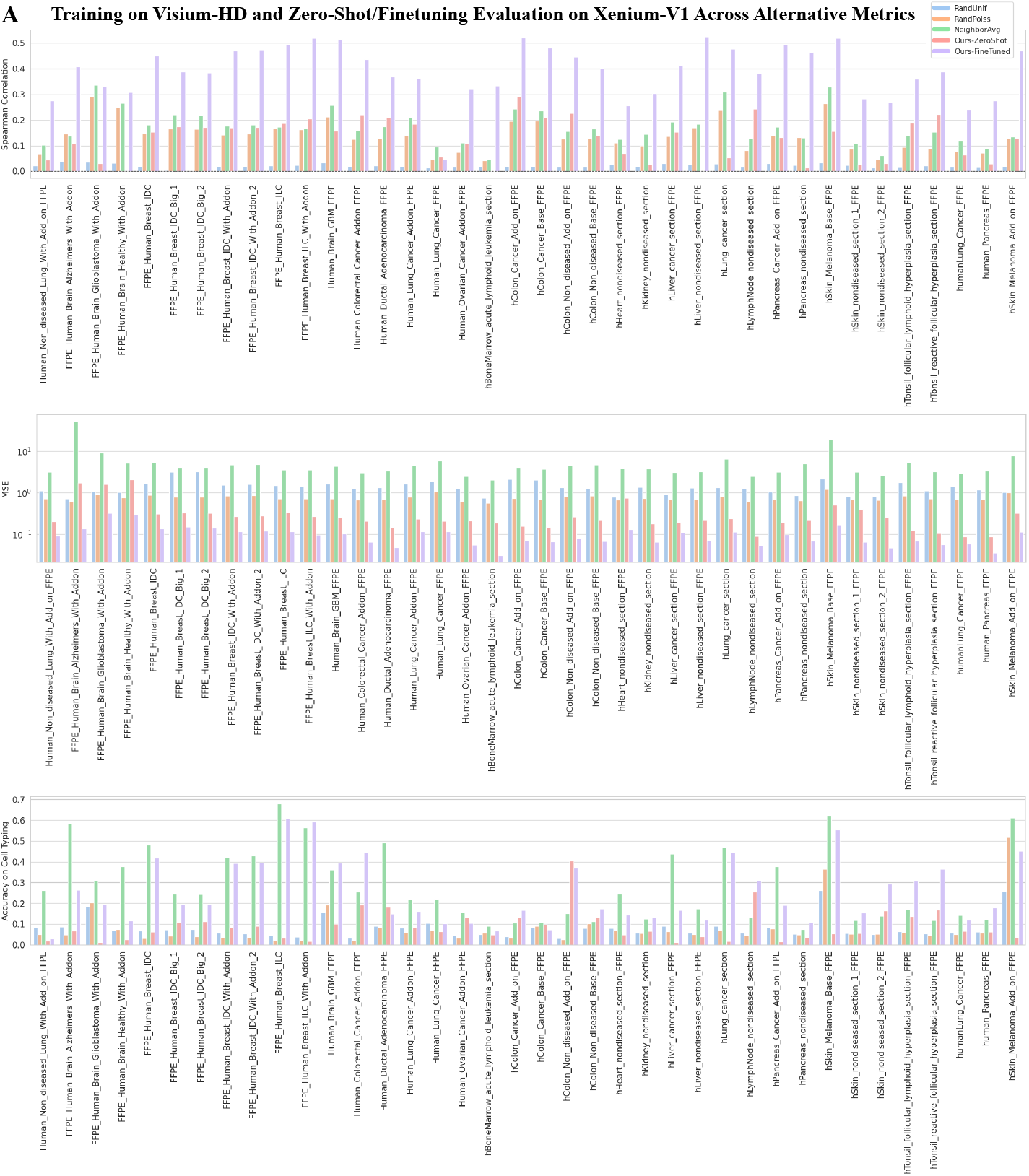
Cross-platform evaluation on the Xenium-V1 dataset. (A) Training on Visium-HD and zero-shot/finetuning evaluation on Xenium-V1. (B) Training on Visium-Prime and zero-shot evaluation on Xenium-Prime.

**Fig. C7].**
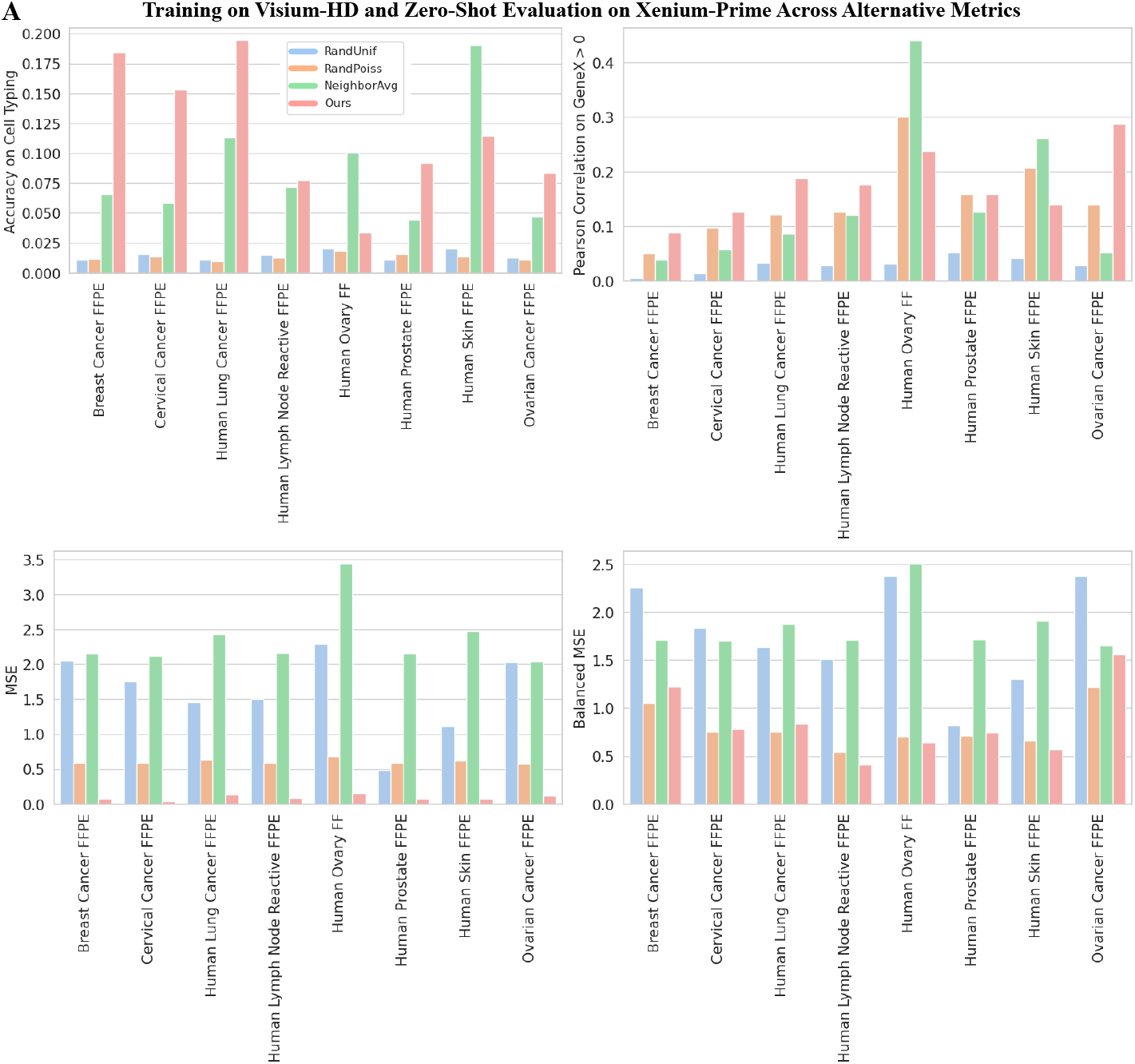
Cross-platform evaluation on the Xenium-Prime dataset. (A) Training on Visium-HD and zero-shot/finetuning evaluation on Xenium-V1. (B) Training on Visium-Prime and zero-shot evaluation on Xenium-Prime.

**Fig. C8].**
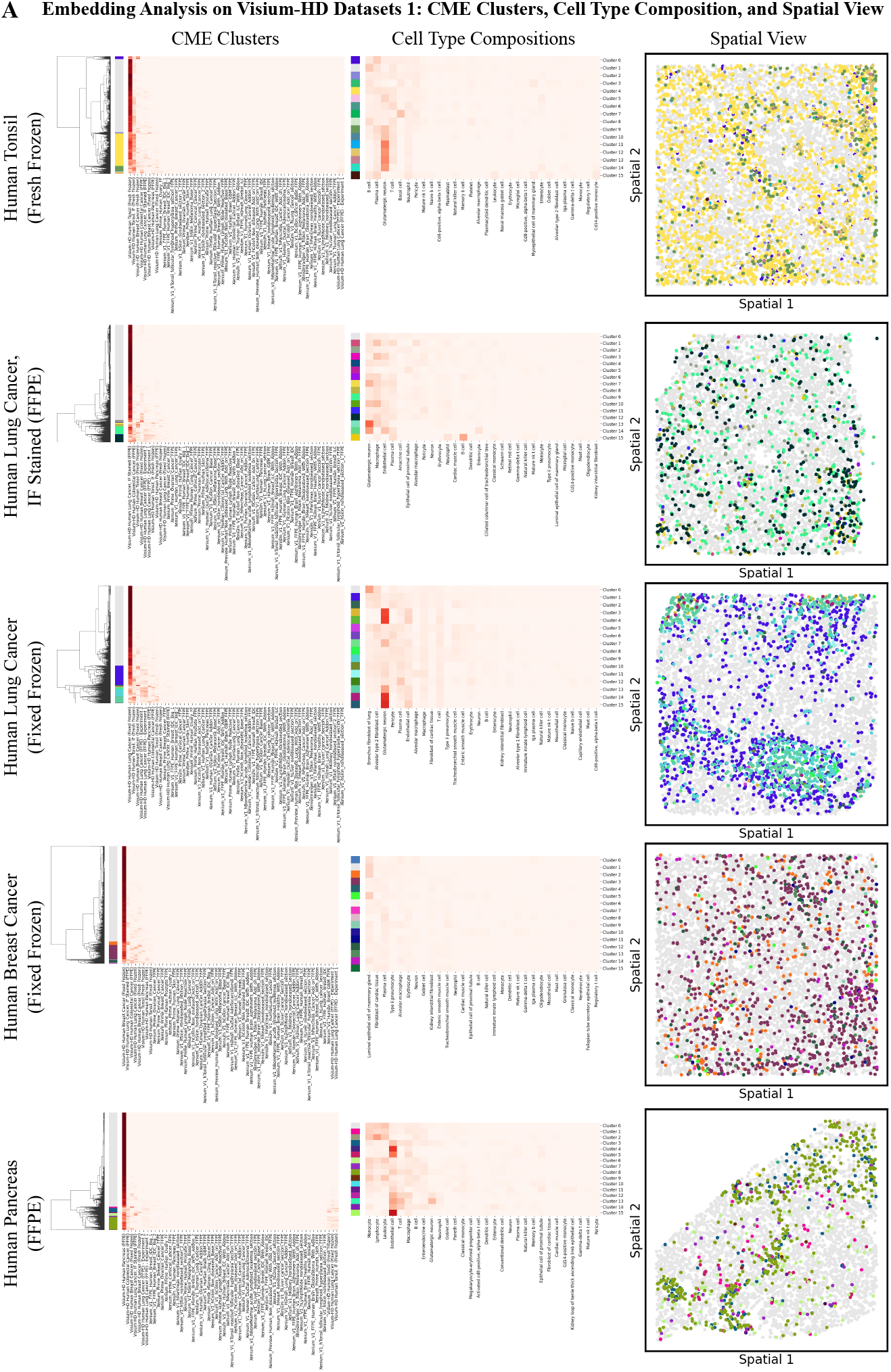
Embedding analysis on other five samples in the Visium-HD dataset. (A) The KNN profile for each CME without averaging, the hierarchical clustering, the cell type composition of each cluster, and the spatial distribution of each CME cluster. The legend is shared with Fig. 3 panel (B).

**Fig. C9].**
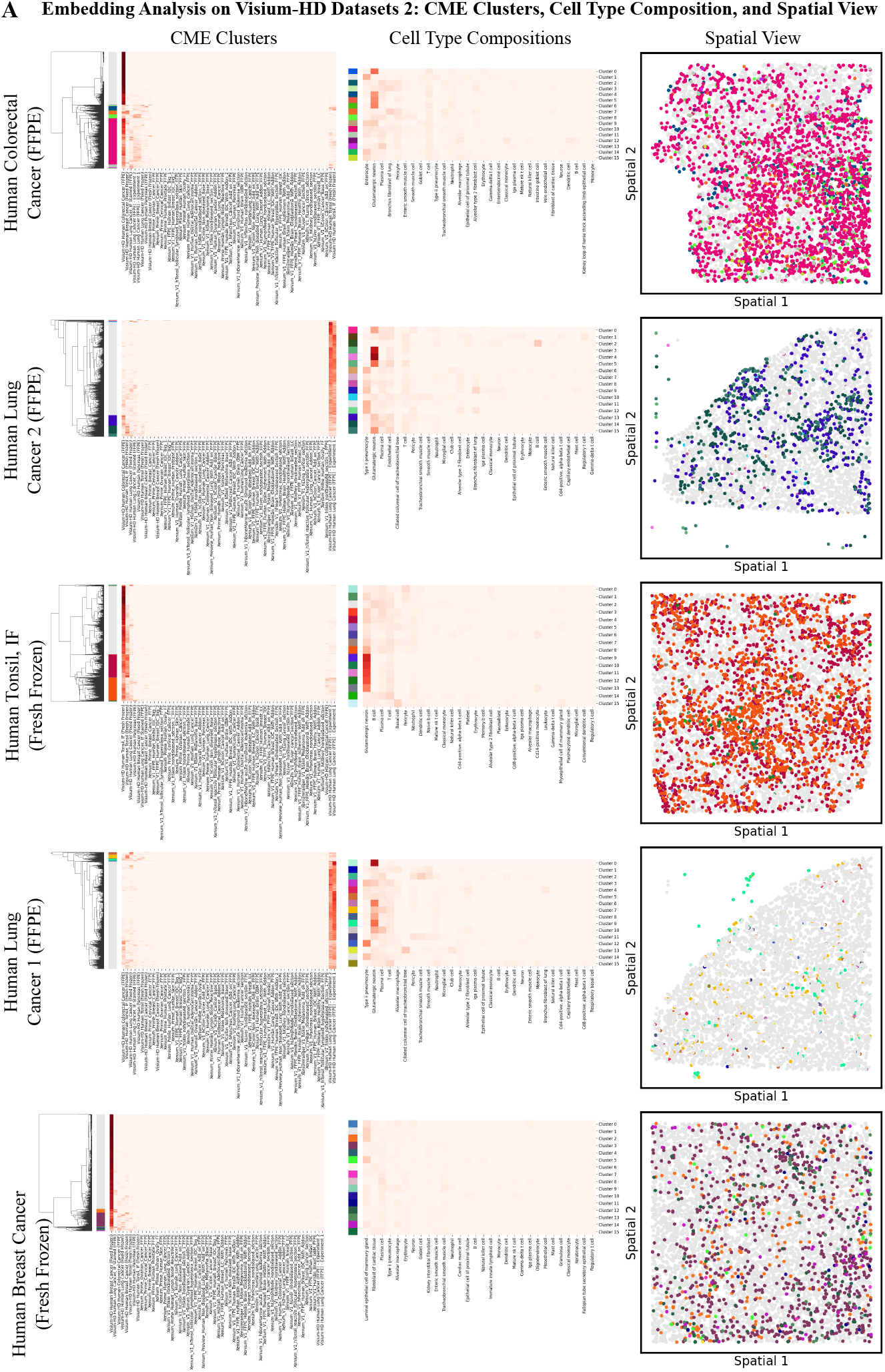
Embedding analysis on other four samples in the Visium-HD dataset. (A) The KNN profile for each CME without averaging, the hierarchical clustering, the cell type composition of each cluster, and the spatial distribution of each CME cluster. The legend is shared with Fig. 3 panel (B).

**Fig. C10].**
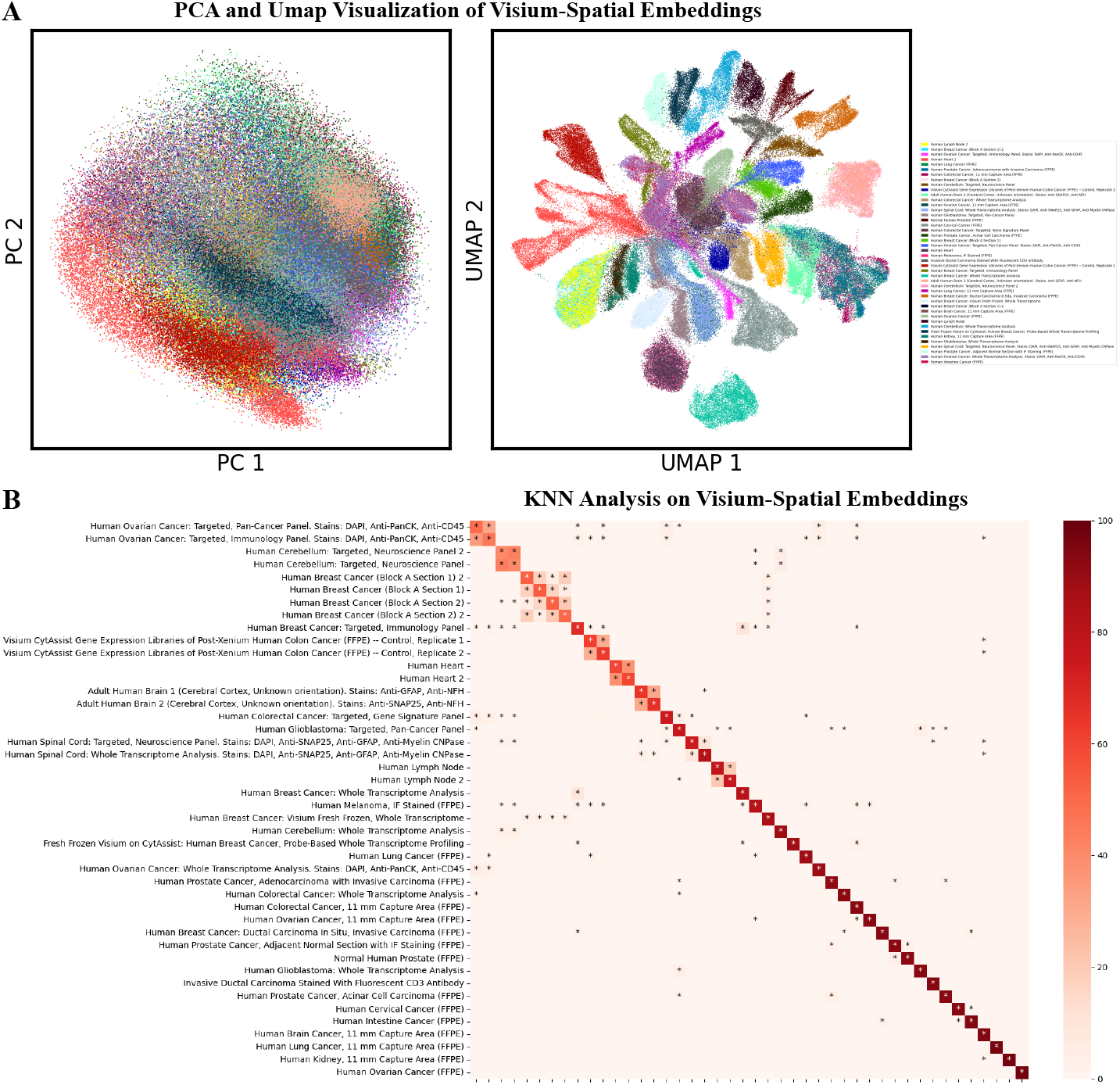
CME embedding with CI-FM on the Visium-Spatial Dataset. (A) Lower-dimensional visualization in PCA and UMAP space. (B) The fraction (%) of the 100 nearest neighbors queried with CI-FM embeddings across different samples. The value is averaged across all CME embeddings of the entire sample. ^*^ denotes a value greater than 1.

**Fig. C11].**
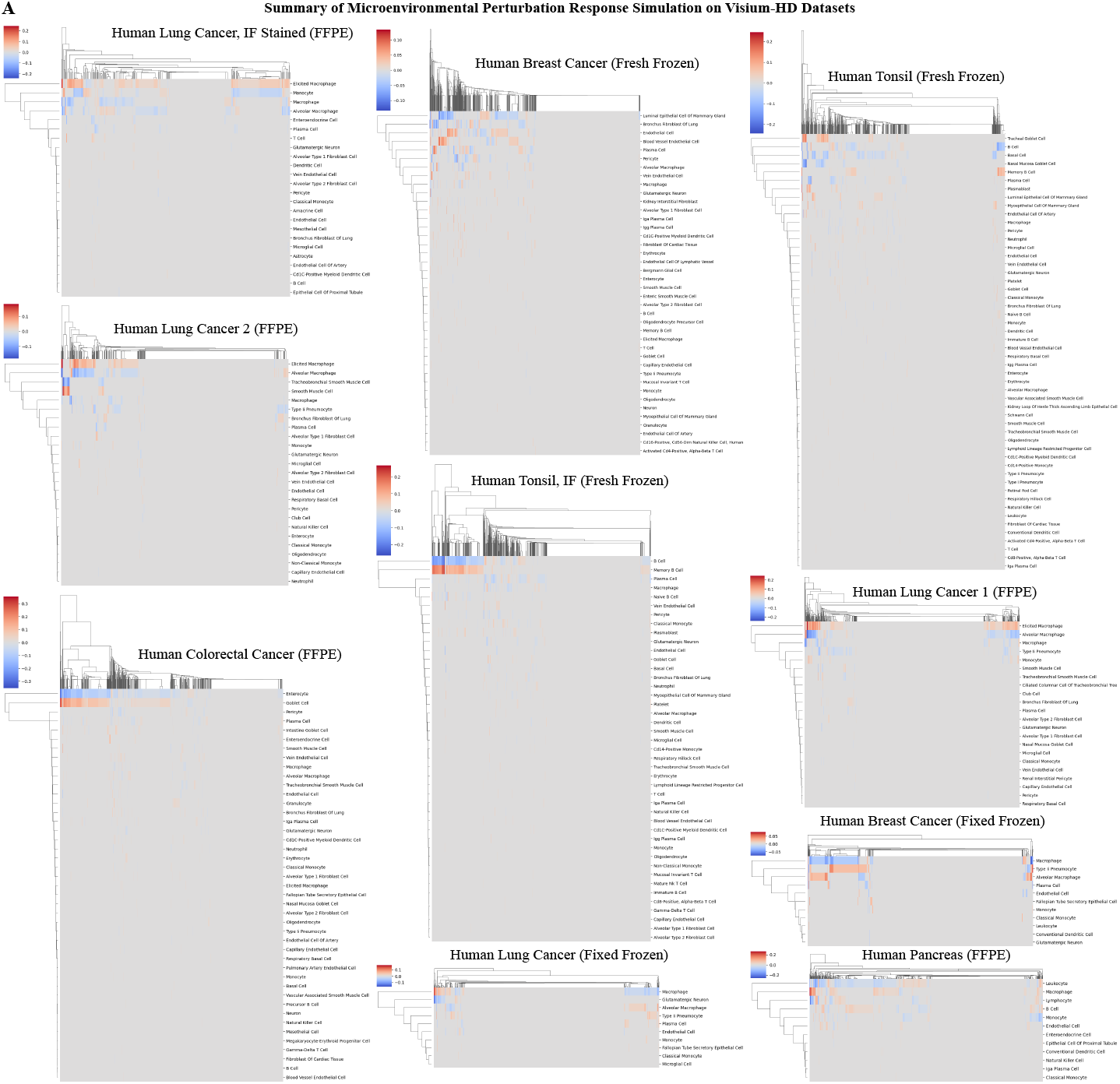
CME response simulation to perturbation with CI-FM. (B) Summary of the cell state change in response to the injection of T cells on other nine samples in the Visium-HD dataset.

